# Distinct Structural Connectivity Patterns Associated with Variations in Language Lateralisation

**DOI:** 10.1101/2025.02.04.636415

**Authors:** Ieva Andrulyte, Laure Zago, Gael Jobard, Herve Lemaitre, Francois Rheault, Simon S Keller, Laurent Petit

## Abstract

Hemispheric asymmetries in white matter tracts are proposed key determinants of language lateralisation, yet evidence in healthy individuals remains inconsistent. This suggests that simple tractography techniques might not be sensitive enough to identify language dominance. Significant insights into the functional organization of the human brain may be achieved by considering networks and brain connectivity, providing more information about discrepancies in people with different hemispheric language dominance. In this study, we examined 285 healthy participants compare their structural connectomes at the whole-brain level and determine the networks responsible for the three different functional language lateralisation groups (typical, atypical and strongly atypical). Probabilistic tractography generated whole-brain tractograms, and white matter fibres were filtered according to anatomical Boolean guidelines. Connectivity matrices with nodes corresponding to supramodal sentence areas in the language atlas and edges weighted by fractional anisotropy (FA) were generated to compare the groups using graph theory and network-based statistic (NBS) approaches. We demonstrated that both atypical (bilateral) and strongly atypical (right-lateralised) lateralisation are characterised by heightened interhemispheric temporal connectivity. Post-hoc analyses showed that strongly atypical individuals exhibited increased temporo-frontal connectivity, while atypical individuals had enhanced temporal and frontal connectivity but lacked temporo-frontal connections. These connectivity patterns diverge from traditional models of hemispheric dominance, suggesting a reliance on integrated bilateral networks in atypically lateralised individuals. This reflects distinct neural mechanisms underlying atypical language organisation, departing from developmental trajectory of typical lateralisation and offering insights into cognitive flexibility and clinical applications.

## Introduction

The lateralisation of language function is thought to be influenced by hemispheric asymmetry in white matter (Verhelst et al., 2021; Westerhausen et al., 2006). However, findings on this relationship in healthy populations remain mixed and inconclusive (Andrulyte et al., 2024; Häberling et al., 2013; Karpychev et al., 2022; Westerhausen et al., 2006), with inconsistencies arising in part from methodological diversity and the limitations posed by small and varied sample sizes. For instance, a recent systematic review of studies using an integrated diffusion MRI and fMRI approach to investigate structural and functional differences across laterality groups revealed that only three studies employed a whole-brain approach, with the remainder relying on region-of-interest (ROI) analyses (Andrulyte et al., 2025). ROI approaches can be useful, provided the selection of tracts is grounded in prior research (Zhang et al., 2022). However, around two-thirds of studies have concentrated solely on the arcuate fasciculus (Andrulyte et al., 2025), traditionally seen as a key tract for language processing in the classical Broca-Wernicke-Geschwind model (Broca, 1861; Geschwind, 1970; Wernicke, Carl, 1874). Although the arcuate fasciculus is crucial for language processing, it is not the sole tract involved, and an exclusive focus on it limits our understanding of the broader structural foundation of language dominance (Tremblay & Dick, 2016). Given that language dominance may be influenced by additional, yet underexplored tracts, the classical model no longer sufficiently accounts for contemporary neurobiological perspectives.

Earlier research has primarily focused on the relationship between laterality indices and individual tract measures, such as FA, streamline count, and volume. However, this approach overlooks a broader, more integrated perspective that is increasingly being adopted in cognitive neuroscience (Fedorenko & Thompson-Schill, 2014). From a cognitive theoretical perspective, functions are viewed as interconnected systems within the brain’s structural and functional connectomes, with cognition emerging from the dynamic interactions of neural networks rather than isolated regions (Bressler & Menon, 2010). The *cognitome*—a term introduced to describe the brain’s cognitive architecture - captures the concept of cognition emerging from complex interactions across multiple brain networks, emphasising their interdependence with structural brain organisation (Roger et al., 2020). This network-based view contrasts with traditional models that isolate functions to discrete regions or tracts, offering a more holistic understanding of cognition.

In the late 20th century, scientists began to shift from traditional views of lateralisation to more complex network-based models. Reggia and colleagues (1998) proposed that the brain operates as a dynamic network of interconnected regions, where synaptic connections adapt through experience, leading to dominant activation patterns. Lateralisation, in this framework, occurs when one hemisphere becomes more active due to factors like increased excitability or learning rates, creating a “race” between hemispheres. This specialisation, particularly in the left hemisphere, helps to enhance processing efficiency and prevent delays that could impact language outcomes (Chiarello et al., 2011; Everts et al., 2009). However, this division of labour hypothesis does not explain why about 10% of individuals rely more on the right hemisphere for language or use both hemispheres for language function (Andrulyte et al., 2024). Recent studies have increasingly examined language lateralisation through a network perspective. For example, fMRI research has shown that atypical individuals—those with bilateral or right-lateralised language—exhibit greater rightward lateralisation within the default mode network, compared to left-lateralised individuals (Labache et al., 2023), suggesting that language lateralisation is dependent on multiple regions working simultaneously. These findings resonate with the previous review of Price (2012) which highlighted that language processing involves over 30 distinct regions, as revealed by PET and fMRI studies, reinforcing the need to shift from traditional ROI-based approaches to more integrated, network-based models (Astrakas, 2023).

To date, only one study has investigated the relationship between structural white matter networks and functional language dominance, revealing higher nodal degrees in the right middle temporal gyrus and posterior cingulate cortex—regions typically associated with the default mode network (Zahnert et al., 2023). However, this study focused solely on nodal properties and their contribution to network organisation, without examining individual subnetworks. Given that the network-based approach has been effective in deciphering white matter networks associated with hemispheric asymmetries (Sun et al., 2017) and different languages (García-Pentón et al., 2014; Wei et al., 2023, 2024), it may also offer a viable method for studying structural differences among individuals with a left-lateralised, right-lateralised, or bilateral language function. The present study aimed to examine structural connectivity to delineate differences across language laterality groups. Diffusion MRI tractography was parcellated into functional cortical regions defined by the SENSAAS atlas (Labache et al., 2019), which includes areas known responsible for language production. The resulting structural connectome was then analysed in a twofold approach: to calculate graph theory measures for individual nodes (Bassett & Sporns, 2017) and to identify subnetworks associated with each laterality group (Zalesky et al., 2010). We hypothesised that (1) distinct white matter networks would be associated with left-, right-, and bilateral language lateralisation; and (2) atypically lateralised individuals would exhibit increased connectivity within right hemisphere regions.

## Methods

### Participants

For this study, we utilised a BIL&GIN dataset (Mazoyer et al., 2016), which is a multimodal imaging database specifically created to investigate hemispheric brain asymmetries in healthy individuals with no history of brain abnormalities. Ethical approval for the study was granted by the Basse-Normandie local Ethics Committee. Participants were selected based on the availability of both sentence production fMRI data and diffusion MRI data, resulting in a final sample of 285 healthy individuals, aged between 18 and 58 years.

### MRI data acquisition

The BIL&GIN data were acquired using a 3T Philips Achieva scanner. High-resolution 3D T1-weighted images were obtained using a 3D-FFE-TFE sequence (TR, 20 ms; TE, 4.6 ms; flip angle, 10 degrees; inversion time, 800 ms; turbo field echo factor, 65; Sense factor, 2; field of view, 256 × 256 × 180 mm; isotropic voxel, 1 × 1 × 1 − mm3). Functional MRI data were obtained with T2*-weighted echo-planar imaging (TR = 2 s, TE = 35 ms, flip angle = 80) across 31 slices, with an isotropic voxel size of 3.75 mm^3^. The sentence production task involved three runs of 192 volumes each (Mazoyer et al., 2016). Diffusion MRI used a spin-echo echo-planar sequence, with a b0 map and 21 diffusion directions (b = 1000 s/mm^2^) acquired twice with reversed polarities. Imaging covered 70 slices from the cerebellum to the vertex (AC-PC plane), using TR = 8500 ms, TE = 81 ms, and a voxel size of 2 mm^3^. For details about the robustness of the diffusion MRI data, see Chenot et al. (2019).

### fMRI preprocessing

Task fMRI data were analysed using Statistical Parametric Mapping (SPM5) software and custom MATLAB routines. For each of the three slow fMRI runs (TR = 2.0 s, voxel size = 3.75 mm^3^), data were corrected for slice timing and motion, with movement-related parameters regressed from the T2*-EPI time series. Scans were aligned to structural images, and T1-weighted volumes segmented into grey matter, white matter, and cerebrospinal fluid. The T2*-EPI scans were normalised to standard stereotaxic space (2 mm^3^ resolution) using a trilinear interpolation. Time series were high-pass filtered with a 159-second cut-off (Joliot et al., 2016).

### Language production task and laterality index

The study used the Word-List production task. During task-based fMRI, BOLD signal variations were measured while participants performed the Word-List production task, which involved generating months of the year to minimise cognitive load on language processing and focus on phonetics and prosody. Each trial began with a 1-second scrambled image, followed by a fixation period where participants completed the task and pressed a response pad. Production tasks lasted 8-14 seconds, while other tasks lasted 6-10 seconds. The total fMRI run was 6 minutes and 24 seconds (Mazoyer et al., 2014). Response times were recorded across ten trials, with participants responding using their right hand.

During the sentence production task, participants were shown line drawings for one second, taken from *Le Petit Nicolas*, followed by a central fixation crosshair. They were asked to covertly generate a sentence that described the scene, starting with a subject and complement, followed by a verb and an additional complement indicating either place or manner (for example, “The little Nicolas… in the street…” or “The gentleman… with happiness…”). In the reference word production task, scrambled versions of these drawings served as stimuli, prompting participants to covertly recite ordered lists, such as the months of the year. Following the sentence generation, participants engaged in a low-level task that required them to detect when the central cross changed into a square, pressing a button upon noticing this transformation. This secondary task was designed to last at least half of the overall trial duration, helping control for any motor responses. Moreover, a 12-second fixation cross was presented before and after the first and last trials to ensure that participants maintained their focus (Labache et al., 2019).

The laterality index (LI) for language production, including its computation and categorisation, was directly adopted from prior analyses by Mazoyer et al. (2014). These analyses, based on “SENT minus LIST” individual t-maps and processed with the LI toolbox (Wilke & Schmithorst, 2006), used a bootstrap algorithm with t = 0, positive t-values, and sample sizes of 5 to 1,000 voxels (resampling ratio k = 0.25). LI values ranged from −100 (right-lateralised) to +100 (left-lateralised), with Gaussian modelling and the corrected Akaike Information Criterion (AICc) identifying three categories: “typical” (LI > 18), “atypical” (−50 < LI ≤ 18), and “strongly atypical” (LI < −50).

### Diffusion MRI processing

Diffusion MRI data processings, including whole-brain probabilistic tractography, were performed using TractoFlow (Theaud et al., 2020), which employs both classical local tracking and Particle Filter Tracking (PFT)(Girard et al., 2014). Default parameters included a step size of 0.5 mm and a maximum angular deviation of 20 degrees. Seeding was conducted across the white matter with 10 seeds per voxel for PFT and 5 seeds per voxel for local tracking, utilising spherical harmonics of order 8. TractoFlow includes a process in which the participant’s T1-weighted image was normalized to the MNI space using ANTS (Avants et al., 2009). Therefore, the affine transformation and nonlinear deformation were applied to warp the whole-brain tractogram to the MNI space.

Anatomically implausible tracts were eliminated from the whole-brain tractogram using the ExTractorflow pipeline, which employs anatomical Boolean rules to extract streamlines that conform to specific anatomical guidelines (Petit et al., 2023). This method is associated with improved reproducibility and precision compared to conventional diffusion MRI techniques, allowing for more accurate identification of anatomically plausible connections through region-of-interest filtering.

We first used SCILPY tools (https://github.com/scilus/scilpy) to segment each whole-brain tractogram into a connectivity matrix using the AICHA parcellation (Joliot et al., 2015). These matrices were thus built between 384 brain regions (192 per hemisphere) by measuring the characteristics of tracts linking each pair of regions, where the nodes represented the parcellated areas, and the edges were weighted by tract length, streamline count, or fractional anisotropy values (FA, for NBS analysis). The initial AICHA-based connectivity matrices were then refined to isolate the 64 supramodal sentence areas, as defined by the SENSAAS language atlas (Labache et al., 2019) (Figure 1).

**Figure 1.**
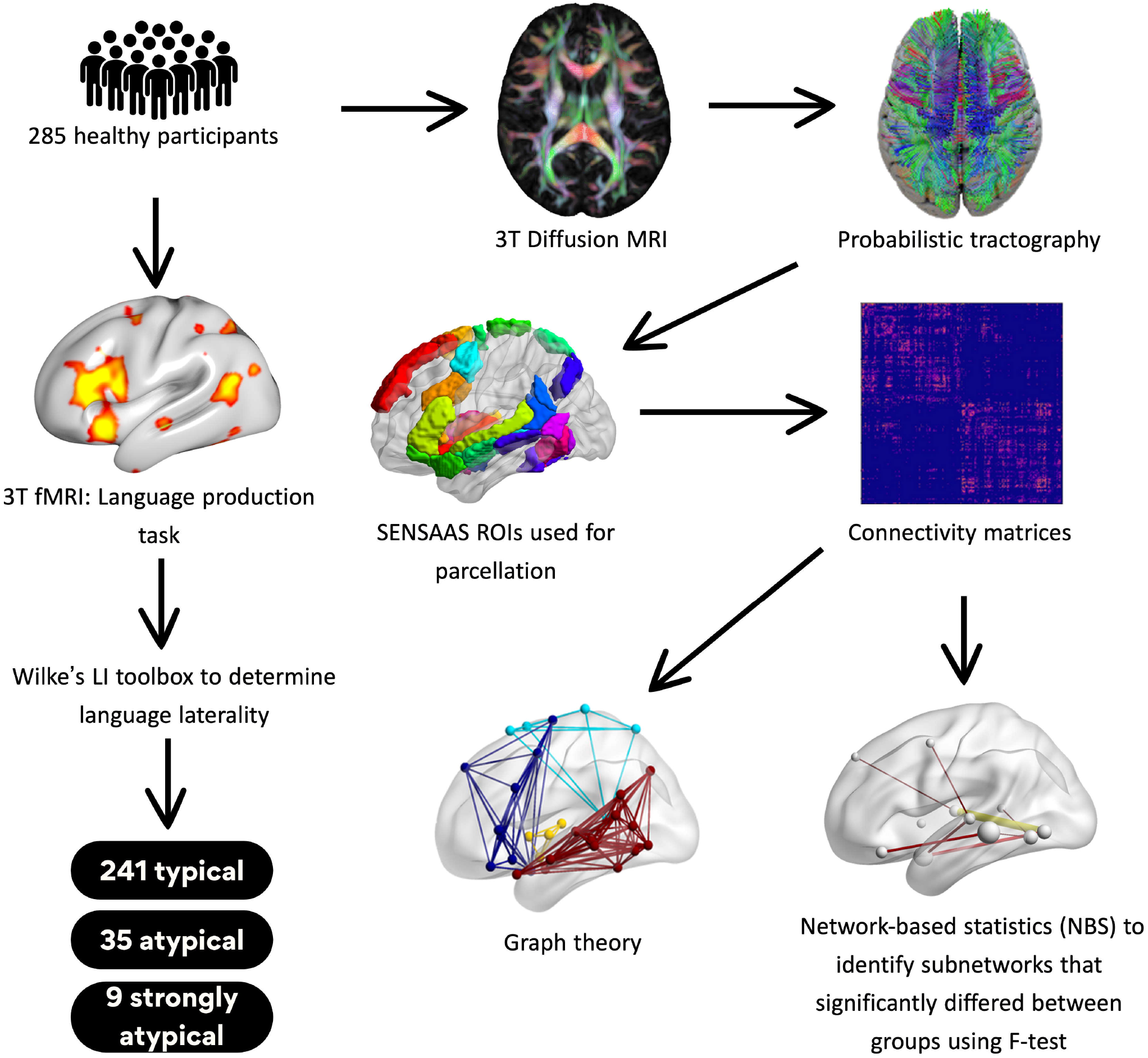
Overview of methodology. This study analysed 285 healthy participants who underwent both fMRI and diffusion MRI scanning. A language production task during 3T fMRI was used to compute language laterality indices via Wilke’s LI toolbox, categorising participants into three groups: 241 with typical lateralisation (LI > 18), 35 with atypical lateralisation (18 > LI > -50), and 9 with strong atypical lateralisation (LI < -50). For diffusion MRI, probabilistic tractography was performed with TractoFlow to produce whole-brain tractograms. Anatomically plausible streamlines were extracted using ExTractorFlow, followed by parcellation with Scilpy to generate connectivity matrices using the AICHA atlas. Subsequent analyses focused on the SENSAAS-defined regions within each hemisphere, measuring global efficiency, nodal strength, local efficiency, path length, and clustering. Group differences across these graph theory metrics were examined using ANOVA and Kruskal-Wallis tests. Additionally, network-based statistics (NBS) were applied to identify subnetworks that significantly differed among the laterality groups.

### Statistical analysis of intra- and interhemispheric connectivity sums

The SENSAAS-based connectivity matrices were used to evaluate both intrahemispheric and interhemispheric connectivity within-subjects across three laterality groups (atypical, strongly atypical, and typical) using connectivity matrices. For each individual, left intrahemispheric connectivity was calculated by summing all connections between left hemisphere nodes, and inversely for the right hemisphere. Interhemispheric connectivity was computed by summing the values of the connections between left and right hemisphere nodes.

Prior to analysis, outliers were identified and excluded based on Mahalanobis distance calculated from the intrahemispheric and interhemispheric connectivity values. Observations exceeding the threshold for multivariate outliers (set at the 97.5% confidence level, corresponding to a chi-squared distribution with degrees of freedom equal to the number of dependent variables) were removed. The resulting intrahemispheric and interhemispheric connectivity values were then analysed between-subjects using multivariate analysis of variance (MANOVA) to assess group differences, with age, sex, and handedness as covariates. Following MANOVA, univariate ANCOVA models were used to investigate the group differences for each connectivity measure: intrahemispheric left FA sum, intrahemispheric right FA sum, and interhemispheric FA sum. Post-hoc pairwise comparisons using Tukey’s test were conducted only for significant ANCOVA results to identify specific group differences. The statistical significance of differences between groups was assessed at the p < 0.05 level.

### Graph theory analysis

The SCILPY tools (https://github.com/scilus/scilpy) were also employed to derive global and local network connectivity metrics from SENSAAS-based connectivity matrices weighted by both streamline count and length for each participant. The analysis focused on computing global efficiency as well as nodal strength, characteristic path length, local efficiency, and clustering coefficient for each node in the SENSAAS atlas (Labache et al., 2019). The selection of these metrics was informed by prior studies indicating their relevance for investigating hemispheric asymmetries (Caeyenberghs & Leemans, 2014; Sha et al., 2022; Y. Zhang et al., 2019).

To compare differences between laterality groups, an analysis of variance (ANOVA) was performed on the global efficiency network metric. This analysis included covariates such as sex, handedness, and age directly within the model. For regional properties where the data did not follow a normal distribution, a Kruskal-Wallis test was used instead. In this case, the network metrics were residualised to account for the effects of covariates, effectively removing any linear associations with sex, handedness, and age before the analysis. This method allowed for the evaluation of the main effects of laterality group on the adjusted network metrics (Figure 1).

### Network-based statistics (NBS) analysis

We used the network-based statistic (NBS) method to identify significant subnetworks comprising differences between laterality groups, as outlined by Zalesky et al. (2010). In essence, NBS is a nonparametric statistical technique specifically designed for analysing large networks, effectively addressing the multiple-comparisons problem that often arises in graph analyses.

To explore differences among typical, atypical, and strongly atypical laterality groups, ANOVA design was employed, utilising FA connectivity matrices. In this analysis, age, sex, and handedness were incorporated as nuisance covariates to control for potential confounding factors. The initial step in the NBS approach involved performing statistical tests on each edge independently, using general linear modelling (GLM) with a threshold F score set at > 4.5, which resulted in the identification of connected pairs.

Following this, NBS examined potential connectivity based on the significant edges and calculated family-wise error rate (FWER)-corrected p-values through permutation testing. This involved assessing connectivity for each permutation alongside the maximal component size, with 5000 permutations carried out at a significance level of p < 0.05. Statistical significance was determined based on an empirical calculation of the null distribution of maximal component sizes across these permutations, where the number of edges in each component represented its size. If a component exhibited a p-value of 0.05 or lower following family-wise correction, it was reported.

The t-test was conducted in MATLAB and applied to FA values of the significant connections identified during the ANOVA analysis. These connections represented edges within the FA connectivity matrices that showed significant differences among the typical, atypical, and strongly atypical laterality groups. Specifically, the comparisons involved typical vs. atypical, atypical vs. strongly atypical, and strongly atypical vs. typical. By focusing on the FA values of these significant connections, the t-test helped assess the directionality of the group differences, indicating whether the FA values were higher or lower for specific laterality groups compared to others. The results were corrected for multiple comparisons using the false discovery rate (FDR) approach to ensure statistical robustness.

## Results

### Demographic characteristics

The language production task identified 241 participants as left-lateralised, 9 as right-lateralised, and 35 as showing bilateral language lateralisation (see Table 1). The mean age of the participants was 25.7 years (SD ± 6.5), with no significant differences in age observed between the lateralisation groups (F-test, p = 0.3). Female participants comprised 48% of the total sample (n = 138), and the distribution of gender did not differ significantly across the lateralisation groups (chi-square, p = 0.33). Handedness was equally distributed, with 48% of participants being right-handed and 52% left-handed. Notably, all participants exhibiting a right-lateralised language pattern (n = 9) were left-handed. The ANOVA showed significant differences in handedness between the lateralisation groups (p = 0.002).

**Table 1.** Demographic information of BIL&GIN data.

|  | *All participants* | *LLD* | *RLD* | *BLR* |
| --- | --- | --- | --- | --- |
| *N, %* | 285 | 241 | 9 | 35 |
| *Mean age (+-SD)* | 25.7(6.5) | 25.5(6.2) | 25.9(6.5) | 27.3(8.2) |
| *Sex (number of females, %)* | 138 (48%) | 118 (49%) | 6 (67%) | 14 (40%) |
| *Handedness Edinburgh Score (number of right-handed people, %)* | 136 (48%) | 123 (51%) | 0 (0%) | 13 (37%) |

LLD -Left language dominance; RLD - Right language dominance; BLR = Bilateral language representation; SD = standard deviation

### Global connectivity strength

A total of six outliers were identified and excluded from the MANOVA analysis. These outliers included n=4 participants from the Typical group, n=1 from the Atypical group, and n=1 from the Strongly Atypical group. The MANOVA revealed no statistically significant effect of laterality group on the combined connectivity measures (Pillai’s Trace = 0.0406, *F* (6, 544) = 1.8791, *p* = 0.082). However, significant effects were observed for the covariates age (p=0.048) and sex (p<0.0001). Handedness did not show a significant effect on connectivity (p=0.292).

### Graph theory

The graph theory results revealed no significant association between the laterality groups with respect to global efficiency (F_2_ = 1.99, p = 0.14). However, significant differences were observed in the covariates: male participants exhibited significantly higher global efficiency than females (F_1_ = 7.25, p = 0.008). Additionally, a significant positive relationship was found between age and global efficiency (F_1_ = 8.85, p = 0.003). Node-based graph theory analyses showed no significant associations after correcting for multiple comparisons (pFDR<0.05) (Supplementary Figure 1).

### NBS analysis

The F-test identified a single connected network of 29 nodes and 30 edges (Figure 2), significantly associated with lateralisation (p < 0.05, F > 4.5, FWER-corrected) (for node abbreviations see Table 2). Upon visual inspection, we observed that the significant edges predominantly connected bilateral temporal and frontal regions. The nodes involved were found in the frontal (superior frontal gyrus [F1_2], inferior frontal sulcus [F2_2], inferior frontal gyrus [F3O1], precentral sulcus [prec3, prec4]), temporal (inferior occipital gyrus [O3_1], superior temporal sulcus [STS3], superior temporal gyrus [T1_4], middle temporal gyrus [T2_3, T2_4], inferior temporal gyrus [T3_4]), subcortical (amygdala [AMYG], thalamus [THA4], putamen [PUT2_L]), insular (anterior insula [INSa1, INSa2, INSa3]), and internal surface areas (posterior cingulate gyrus [CINGp3], paracentral lobule [pCENT4], precuneus [PRECU6], supplementary motor area [SMA3]).

**Table 2.**
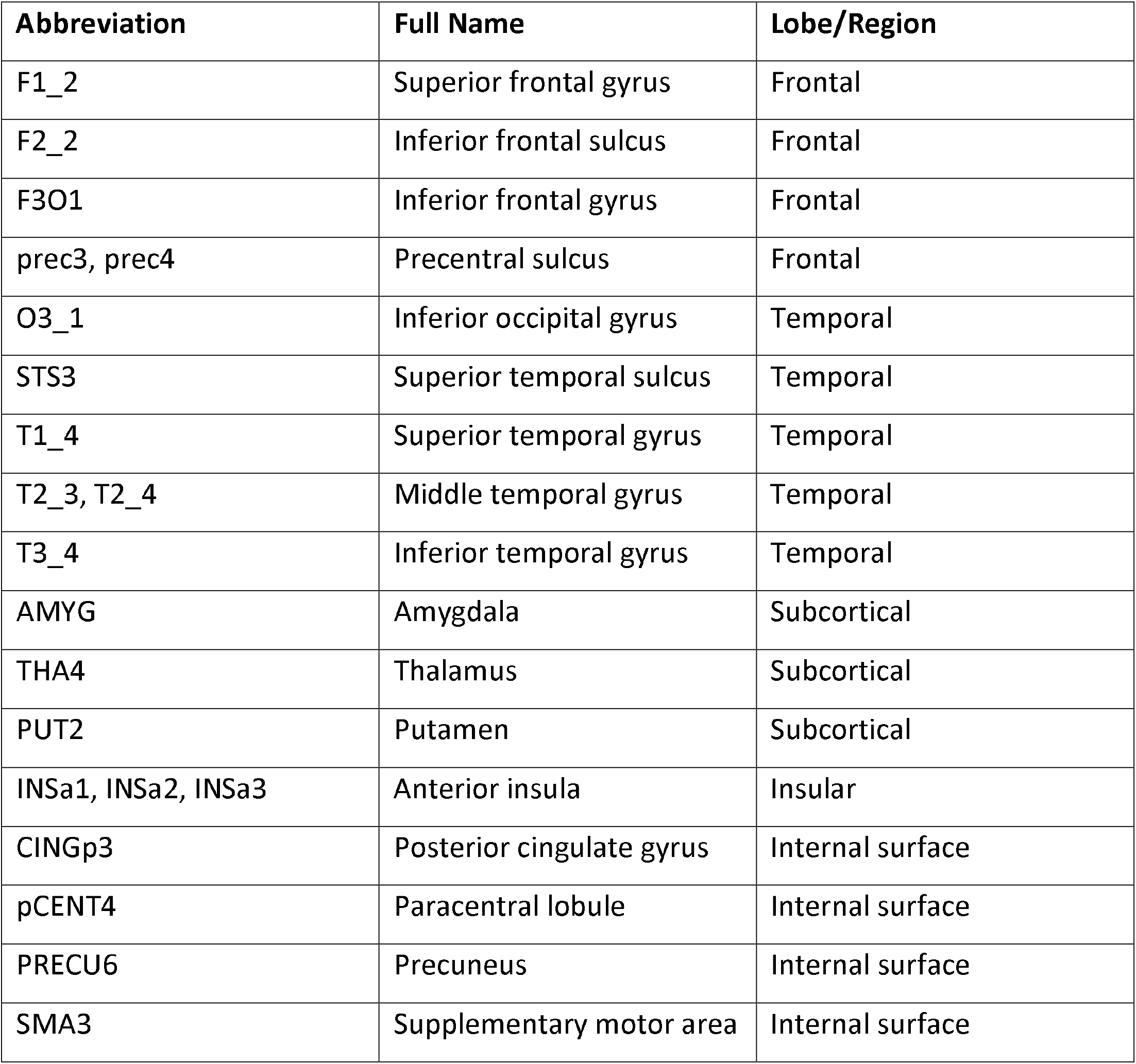
Node abbreviations and anatomical regions.

| Abbreviation | Full Name | Lobe/Region |
| --- | --- | --- |
| F1_2 | Superior frontal gyrus | Frontal |
| F2_2 | Inferior frontal sulcus | Frontal |
| F3O1 | Inferior frontal gyrus | Frontal |
| prec3, prec4 | Precentral sulcus | Frontal |
| O3_1 | Inferior occipital gyrus | Temporal |
| STS3 | Superior temporal sulcus | Temporal |
| T1_4 | Superior temporal gyrus | Temporal |
| T2_3, T2_4 | Middle temporal gyrus | Temporal |
| T3_4 | Inferior temporal gyrus | Temporal |
| AMYG | Amygdala | Subcortical |
| THA4 | Thalamus | Subcortical |
| PUT2 | Putamen | Subcortical |
| INSa1, INSa2, INSa3 | Anterior insula | Insular |
| CINGp3 | Posterior cingulate gyrus | Internal surface |
| pCENT4 | Paracentral lobule | Internal surface |
| PRECU6 | Precuneus | Internal surface |
| SMA3 | Supplementary motor area | Internal surface |

**Figure 2.**
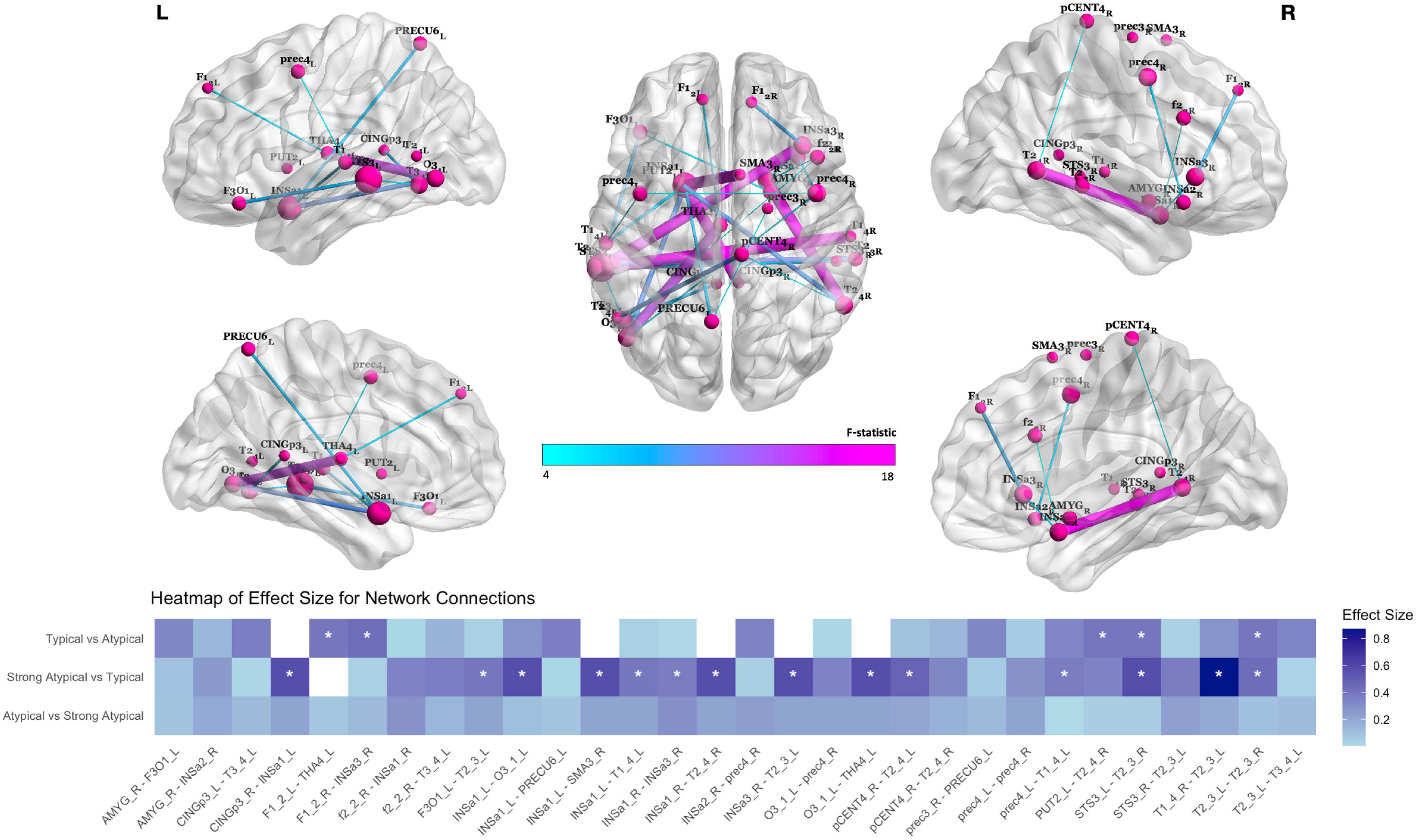
Intergroup differences in structural connectivity strength across language laterality groups. Structural connectivity network depicts connections associated with language production lateralisation across the three laterality groups, as determined by ANOVA (p<0.05, FWE-corrected). This network consists of 29 nodes and 30 edges, with edge weights based on FA values. Significant differences in connectivity between the groups were identified using NBS method. The colour scale represents the F-statistic values. Heatmap displays effect sizes for each connection based on post-hoc t-tests conducted on all significant connections identified using NBS. Three separate t-tests were performed between each pair of groups (typical vs atypical, strong atypical vs typical, and atypical vs strong atypical) and corrected for false discovery rate (FDR). Asterisks (*) indicate statistically significant differences after FDR correction. A complete list of abbreviations is provided in Table 2. ‘L’ and ‘R’ denote the left and right hemispheres, respectively.

Post-hoc t-tests revealed that five of the 30 connections were significantly associated with atypical lateralisation (as compared to typical lateralisation, Figure 3 left). Most of these connections were observed in the frontal and temporal lobes, with a few in subcortical regions. Both intra- and inter-hemispheric connections were present, and the strongest connection (t > 4) was detected between the right anterior insula (INSa3_R) and the right superior frontal gyrus (F1_2_R) (pFDR = 0.004). The right middle temporal gyrus (T2_3_R) showed the highest nodal degree (n = 2), with significant connections to the left superior temporal sulcus (STS3_L) (pFDR = 0.017) and left middle temporal gyrus (T2_3_L) (pFDR = 0.018). Additional significant connections included the left putamen (PUT2_L) to the right middle temporal gyrus (T2_4_R) (pFDR = 0.017) and the left superior frontal gyrus (F1_2_L) to the left thalamus (THA4_L) (pFDR = 0.017) (Figure 3 left). No significant connections were found in the opposite comparison (typical vs. atypical lateralisation) after correction for multiple comparisons.

**Figure 3.**
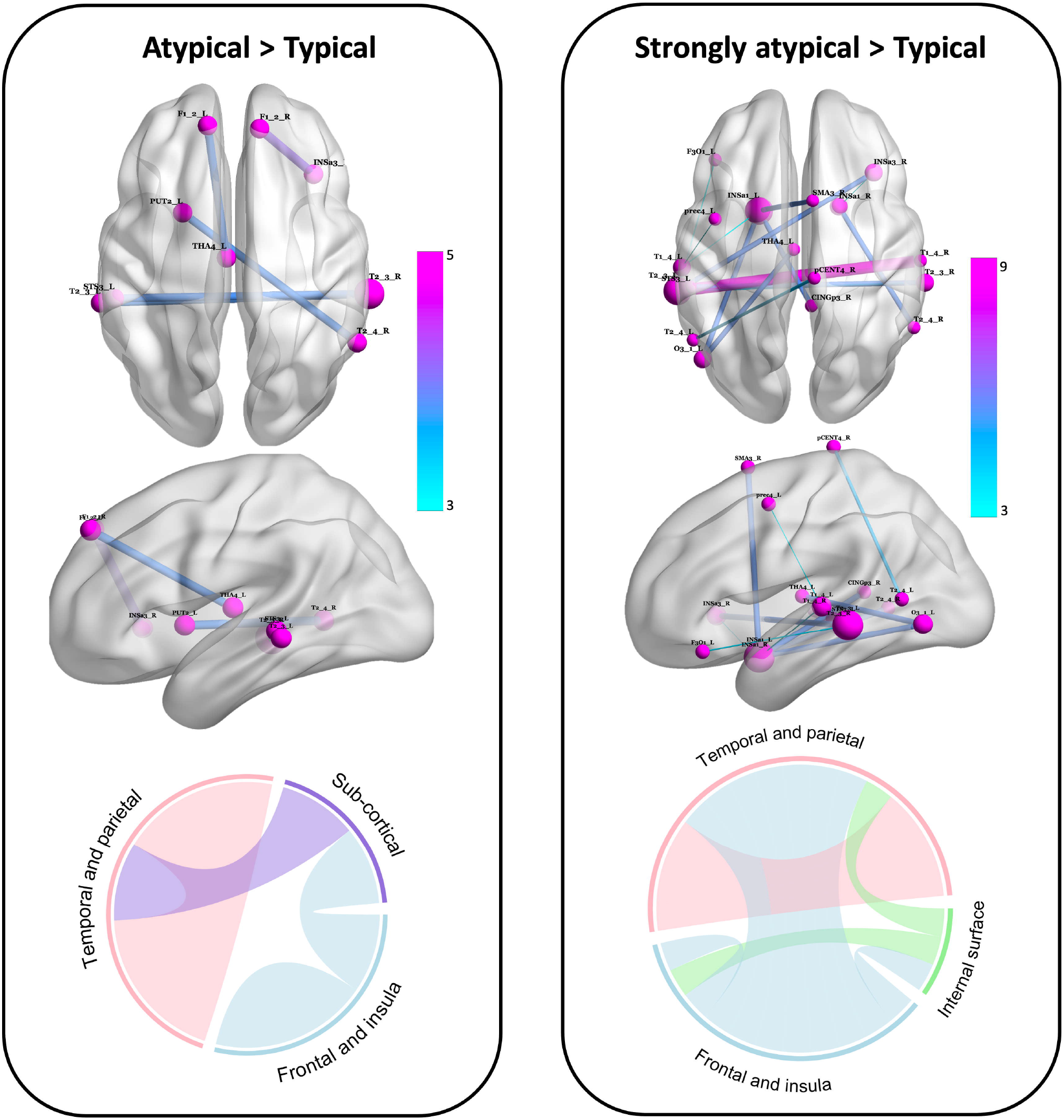
Post-hoc analysis of structural connectivity differences between language laterality groups. **(Left)** Connections showing significantly greater structural connectivity strength in the Atypical group compared to the Typical group. These connections are primarily located within the temporal and parietal lobes, subcortical areas, and the frontal and insula lobes. **(Right)** Connections with significantly greater connectivity strength in the Strongly Atypical group relative to the Typical group. Connections are concentrated between temporal and parietal lobes, the internal surface, and frontal and insula regions. Node sizes again indicate nodal strength, and edge colours represent F-statistic values, with pink for higher values and blue for lower values. The colour bars indicate the range of t-statistic values for each comparison. The Circos plots at the bottom of each panel summarise the main connectivity patterns across four anatomical divisions: temporal and parietal regions, subcortical areas, frontal and insular cortices, and internal surface structures. In these plots, colour-coded segments represent the anatomical divisions, and the widths of these segments reflect the number of nodes and their overall connectivity strength within the respective division. Abbreviations for anatomical regions are detailed in Table 2, with ‘L’ and ‘R’ denoting the left and right hemispheres, respectively.

In individuals with strongly atypical lateralisation (as compared to typical lateralisation, Figure 3 right), the connections were primarily between regions of the insular and temporal lobes, alongside some internal surface regions, displaying a mix of intra- and inter-hemispheric links. The strongest connection (t > 8) was between the right superior temporal gyrus (T1_4_R) and the left middle temporal gyrus (T2_3_L) (pFDR < 0.0001). The left middle temporal gyrus (T2_3_L) demonstrated the highest nodal degree (n = 5), with connections to the left inferior frontal gyrus (F3O1_L) (pFDR = 0.019), right anterior insula (INSa3_R) (pFDR < 0.0001), superior temporal sulcus (STS3_R) (pFDR < 0.001), right superior temporal gyrus (T1_4_R) (pFDR < 0.0001), and right middle temporal gyrus (T2_3_R) (pFDR = 0.003). The left anterior insula (INSa1_L) exhibited the second-highest nodal degree (n = 4), with connections to the left inferior occipital gyrus (O3_1_L) (pFDR < 0.0001), right supplementary motor area (SMA3_R) (pFDR < 0.0001), left superior temporal gyrus (T1_4_L) (pFDR = 0.025), and right cingulate gyrus (CINGp3_R) (pFDR < 0.0001). Further connections included the right anterior insula 1 (INSa1_R) to right anterior insula 3 (INSa3_R) (pFDR = 0.04), the left precentral sulcus (prec4_L) to left superior temporal gyrus (T1_4_L) (pFDR = 0.039), and the left superior temporal sulcus (STS3_L) to right middle temporal gyrus (T2_3_R) (pFDR < 0.0001). No significant connections were found in the opposite comparison (typical vs. strongly atypical lateralisation) after correction for multiple comparisons..

Finally, the comparison between strongly atypical and atypical individuals showed no significant differences after correction for multiple comparisons.

## Discussion

In our study, we found that (1) the NBS approach identified connections in the sentence processing network that significantly distinguish three laterality groups involving bilateral temporal, frontal, and insular cortices; (2) bilateral individuals have greater FA connectivity in bilateral temporal connections and intrahemispheric frontal connections compared to left lateralised individuals; (3) right lateralised individuals also demonstrate increased bilateral temporal lobe FA connectivity, along with enhanced connectivity between frontal/insular and temporal regions; and (4) graph theory analysis revealed no differences (after FDR correction) in global or local connectivity between the groups.

### Increased interhemispheric connectivity in bilateral and right lateralised people

Our findings indicate that individuals with atypical and strongly atypical language lateralisation exhibit increased interhemispheric connectivity, as evidenced by greater FA values in temporal lobe bilaterally, compared to those with typical lateralisation. This supports our previous findings demonstrating a significant association between higher FA values in commissural fibres and a decreased laterality index in semantic comprehension tasks in an independent cohort of healthy individuals (Andrulyte et al., 2024). The only other study exploring the same edge-level connectivity in relation to language lateralisation used resting-state fMRI data from the BIL&GIN dataset and reported that atypical lateralisation is linked to greater interhemispheric connectivity, particularly within the frontotemporal network (Labache et al., 2020). However, the underlying mechanisms and reasons for these differences, particularly in relation to diffusion-based white matter measures, remain unclear.

Traditional models of language lateralisation describe a shift from diffuse activation to left-dominant patterns during development (Friederici et al., 2011; Perani et al., 2011; Szaflarski et al., 2006; Kadis et al., 2011), associated with changes such as callosal shrinkage (Karolis et al., 2019) and maturation of the left arcuate fasciculus (Tak et al., 2016), which supports increased intrahemispheric connectivity between the frontal and temporal lobes (Reynolds et al., 2019; Tzourio-Mazoyer et al., 2017). In contrast, atypical lateralisation may follow an alternative developmental trajectory, with some evidence linking it to musicianship (Villar-Rodríguez, Marin-Marin, et al., 2024). While Villar-Rodríguez et al. (2024) did not investigate developmental mechanisms, they found that atypically lateralised musicians exhibit greater interhemispheric connectivity and larger corpus callosum volume, while non-musicians with atypical lateralisation show underdevelopment of the left arcuate fasciculus. This suggests two potential developmental pathways for language lateralisation.

### Atypical lateralisation extends beyond a mirrored network of left-lateralised individuals

Although the mechanisms underlying the development of atypical language dominance remain poorly understood, Price & Crinion (2005) proposed that atypical dominance arises from inhibition of the non-dominant hemisphere by the dominant hemisphere, resulting in activation of homologous or mirrored regions. Initially based on studies in individuals with aphasia, the model was later extended to healthy populations and appeared supported by evidence at the time. For example, fMRI studies showed that right-dominant individuals exhibited activation patterns in the right hemisphere that mirrored those of left-lateralised individuals in the left hemisphere during language tasks (Bidula et al., 2017; Knecht et al., 2003). Combined task fMRI and whole diffusion MRI studies further supported this model, reporting reduced FA in the left arcuate fasciculus (Ocklenburg et al., 2013) and left superior longitudinal fasciculus (Perlaki et al., 2013) in individuals with atypical lateralisation. This model remained dominant until relatively recently.

Recent evidence challenges this model, with whole-brain diffusion MRI studies reporting no significant association between fixel-based diffusion metrics of the arcuate fasciculus and language lateralisation, despite observing leftward arcuate fasciculus asymmetry in fixel-based metrics that was not directly linked to functional language dominance (Verhelst et al., 2021). This shift is supported by recent fMRI research, which suggests more complex mechanisms underpinning language lateralisation. For example, although Bidula et al. (2017) demonstrated mirrored activation patterns in the right hemisphere of right-lateralised individuals, bilateral individuals exhibited a more diffuse activation pattern, involving regions such as the middle temporal gyrus (MTG) and posterior cingulate cortex. Our findings similarly show bilateral MTG connectivity in both right-lateralised and bilateral individuals compared to typically lateralised individuals, alongside increased connectivity between the right posterior cingulate gyrus and left anterior insula in right-lateralised individuals. Moreover, neither bilateral nor right-hemisphere-dominant individuals showed a clear hemispheric preference but exhibited increased FA in both hemispheres, a finding supported by recent functional MRI studies (Labache et al., 2020, 2023; Villar-Rodríguez, Cano-Melle, et al., 2024). One such study reported an increase in the volume of the corpus callosum in atypically lateralised individuals (Labache et al., 2020). However, this study used the same dataset as in our study, highlighting the need for future research to replicate these findings across different language tasks and further explore the coupling between structural and functional networks. These studies are crucial for understanding the developmental trajectory of atypical language lateralisation, which cannot be explained by a simple inversion of typical leftward hemispheric dominance and highlight the need for more comprehensive models to account for its complexity.

### The role of the left and right temporal lobes in atypical language dominance

According to Hickok and Poeppel’s dual-stream model of language, the ventral tracts are bilateral and play a crucial role in language processing. These tracts connect regions such as the superior temporal sulcus, MTG, and superior temporal gyrus, which are essential for mapping sounds to meanings (Catani & Bambini, 2014; Moritz-Gasser et al., 2013). The bilateral nature of the ventral stream highlights that not all aspects of language processing are strictly lateralised, helping explain the minimal phonemic perception deficits observed after unilateral lesions to the superior temporal gyrus. In contrast, bilateral damage to these regions results in more severe impairments, such as word deafness (Poeppel, 2003). Binder et al. (2024) mapped interhemispheric connections in the temporal lobes, revealing extensive transcallosal projections, particularly involving the MTG bilaterally. This therefore might explain how disruptions to the left MTG—commonly associated with language impairments—may not solely reflect its role in language processing but rather the disturbance of interhemispheric communication, notably via the corpus callosum (Hickok, 2022; Maffei et al., 2017; Northam et al., 2012).

The role of temporal lobe bilaterally in language processing is well established, but the increased interhemispheric connectivity observed in the MTG in atypical individuals (i.e., bilateral or right-lateralised people) raises the question of why this heightened connectivity occurs specifically in these individuals. fMRI studies suggest that children initially exhibit strong interhemispheric temporal connectivity, which transitions to more intrahemispheric frontotemporal connectivity as they mature (Friederici et al., 2011). This developmental trajectory implies that the increased bilateral connectivity in atypical individuals could reflect a disruption in the typical development of language networks, where the shift from interhemispheric to intrahemispheric connectivity does not occur as it does in typical individuals. Longitudinal studies are crucial to explore this developmental trajectory further, as they are the only way to directly assess how these atypical patterns of connectivity evolve over time and their impact on language lateralisation.

### The role of the anterior insula in atypical language dominance

The anterior insula’s role in language processing has been well-documented (Oh et al., 2014); however, its specific relationship with hemispheric language dominance remains underexplored. The present analysis revealed distinct connectivity patterns in individuals with atypical language lateralisation, indicating a stronger engagement of the anterior insula compared to those with typical left-lateralised language dominance. Notably, individuals with right language lateralisation exhibited heightened anterior insula connectivity, characterised by significant bilateral engagement in both intra- and interhemispheric connections. Conversely, the group with bilateral language representation demonstrated the strongest connectivity intrahemispherically within the right hemisphere, particularly between the anterior insula and the superior frontal gyrus (SFG).

These findings are somewhat comparable to a few structural MRI grey matter studies, such as Keller et al. (2011), who reported a rightward insular asymmetry in individuals with right-hemisphere dominance and demonstrated that this asymmetry could predict rightward lateralisation with 90% accuracy. Similar associations between insular asymmetry and functional language dominance have been observed in other studies (Biduła & Króliczak, 2015; Gerrits et al., 2022; Greve et al., 2013; Keller et al., 2018). However, this study adds a novel perspective by highlighting distinct anterior insula connectivity patterns in atypically lateralised individuals. Notably, the heightened bilateral involvement in the right-lateralised group suggests a more flexible and less rigidly organised language network, aligning with Greve et al. (2013), who associated lower left insular asymmetry with atypical language lateralisation.

One plausible explanation for the anterior insula’s increased engagement in atypical individuals lies in the challenges associated with integrating linguistic information, which may be attributed to the diffuse and inefficient nature of language processing in these individuals, in contrast to the more streamlined left-lateralised processing typically observed in those with standard language dominance (Ringo et al., 1994). Recent research indicates that the right anterior insula facilitates cognitive tasks by suppressing the default mode network, thereby improving attentional focus in demanding contexts (Raichle, 2015), such as foreign language processing (Rogenmoser et al., 2023). This suggests that atypically lateralised individuals may rely on the right anterior insula more frequently to manage attentional shifts critical for language comprehension and production. Ferstl et al. (2005) demonstrated right insular activation during narrative comprehension tasks involving global contextual inconsistencies, implicating these regions in managing heightened integration demands. Similarly, Mason and Just (2007) highlighted the right anterior insula’s involvement in resolving interference during sentence comprehension tasks with polysemous words, emphasising its role in suppressing incorrect interpretations. These findings suggest that atypically lateralised individuals, who lack the streamlined hemispheric organisation typically associated with language, may depend on the right anterior insula for inhibitory control and interference resolution during complex linguistic tasks (Bari & Robbins, 2013; Deng et al., 2018). This reliance on the anterior insula may reflect compensatory mechanisms supporting language processing in the absence of typical lateralisation patterns.

### Methodological considerations

In the present study, several limitations warrant careful consideration. Firstly, the atlas used in our analyses was constructed based solely on right-handed individuals (Labache et al., 2019), while our sample included both right- and left-handed participants. While this could suggest potential limitations in capturing the full spectrum of brain regions involved in language processing, particularly those with differential activation patterns in left-handed individuals, the impact on our study is likely minimal. It is well-documented that brain architecture can vary substantially between left- and right-handers on the functional level (Johnstone et al., 2021); however, structural differences are less clear, with only grey matter differences observed (Sha et al., 2021), and no significant differences found in diffusion MRI connectivity (Tomasi & Volkow, 2024). Furthermore, the atlas includes key high-order language regions (Fedorenko et al., 2024), meaning that while the mismatch between handedness in the atlas and our sample may pose a theoretical concern, it is unlikely to have substantially affected the validity of our results.

Second, a common limitation in network-based studies, including ours, is the balance between false positives and false negatives when correcting for multiple comparisons. While uncorrected analyses risk inflating false positives, stringent correction procedures reduce statistical power (Helwegen et al., 2023). To mitigate this, we adopted the NBS approach, which assesses connected components of edges rather than individual edges, thereby enhancing statistical power at a strict significance threshold (Zalesky et al., 2010). The use of cyclic topology-focused approaches like NBS is well-supported, as they improve the reliability of identifying significant connectivity patterns (Baggio et al., 2018). It is important to acknowledge, however, that NBS inherently prioritises topological features, which may reduce specificity for individual edges. This could potentially result in missed focal connectivity patterns, particularly those with lower clustering or less cyclic topology (Helwegen et al., 2023). Understanding this aspect of the methodology helps clarify why certain effects, such as the significance of individual edges across the whole brain, may not be detected. To address this limitation, our post-hoc analyses focused on individual edges within the significant edges found in the network/cluster detected by the F-test NBS. While this step identified large effect sizes (greater than 0.7) for these edges (Figure 2), it is important to note that post-hoc analyses are limited to edges within already significant clusters and do not fully mitigate this issue across the whole brain. Nonetheless, this approach minimised the impact of this limitation on our findings.

Lastly, statistical power was limited by the small sample size of strongly atypical individuals (n=9), despite the overall large sample size (n=285). The rarity of strongly atypical lateralisation (∼4-5%) in the general population (Biduła & Króliczak, 2015) accounts for the small size of this subgroup. While this limitation reduces statistical power, it is important to note that the small number of right-lateralised individuals reflects demographic trends, and there is no feasible solution to overcome this. Despite this, the study represents one of the most extensive investigations into language lateralisation in relation to diffusion MRI measures, with one of the largest sample sizes in the field to date (Andrulyte et al., 2025).

## Conclusion

We investigated connectivity differences across individuals with strongly atypical, atypical, and typically lateralised language processing using graph theoretical and network-based methods. Our findings reveal that individuals with atypical lateralisation exhibit a more diffuse connectivity pattern, characterised by heightened interhemispheric connectivity, particularly in those with strongly atypical lateralisation (i.e. moderately to strongly right lateralised). Additionally, we observed enhanced connectivity within the anterior insula, with increased right intrahemispheric connectivity in bilaterally lateralised individuals and both intra- and interhemispheric activity in right-lateralised individuals, emphasising its role in language lateralisation. These findings contribute to the growing evidence that atypical language lateralisation is linked to distinct connectivity patterns that diverge from typical developmental paths, offering valuable insights into how language processing works in atypical populations. They highlight the need for further research into the functional and clinical implications of these connectivity differences. Longitudinal studies would be particularly useful in tracking how language lateralisation develops in atypical individuals, shedding light on the progression of these connectivity patterns and their impact on language acquisition and cognitive abilities while also helping to identify the factors that influence the development of atypical lateralisation.

