## Supplemental Fig. 1 for "Distinct Structural Connectivity Patterns Associated with Variations in Language Lateralisation"

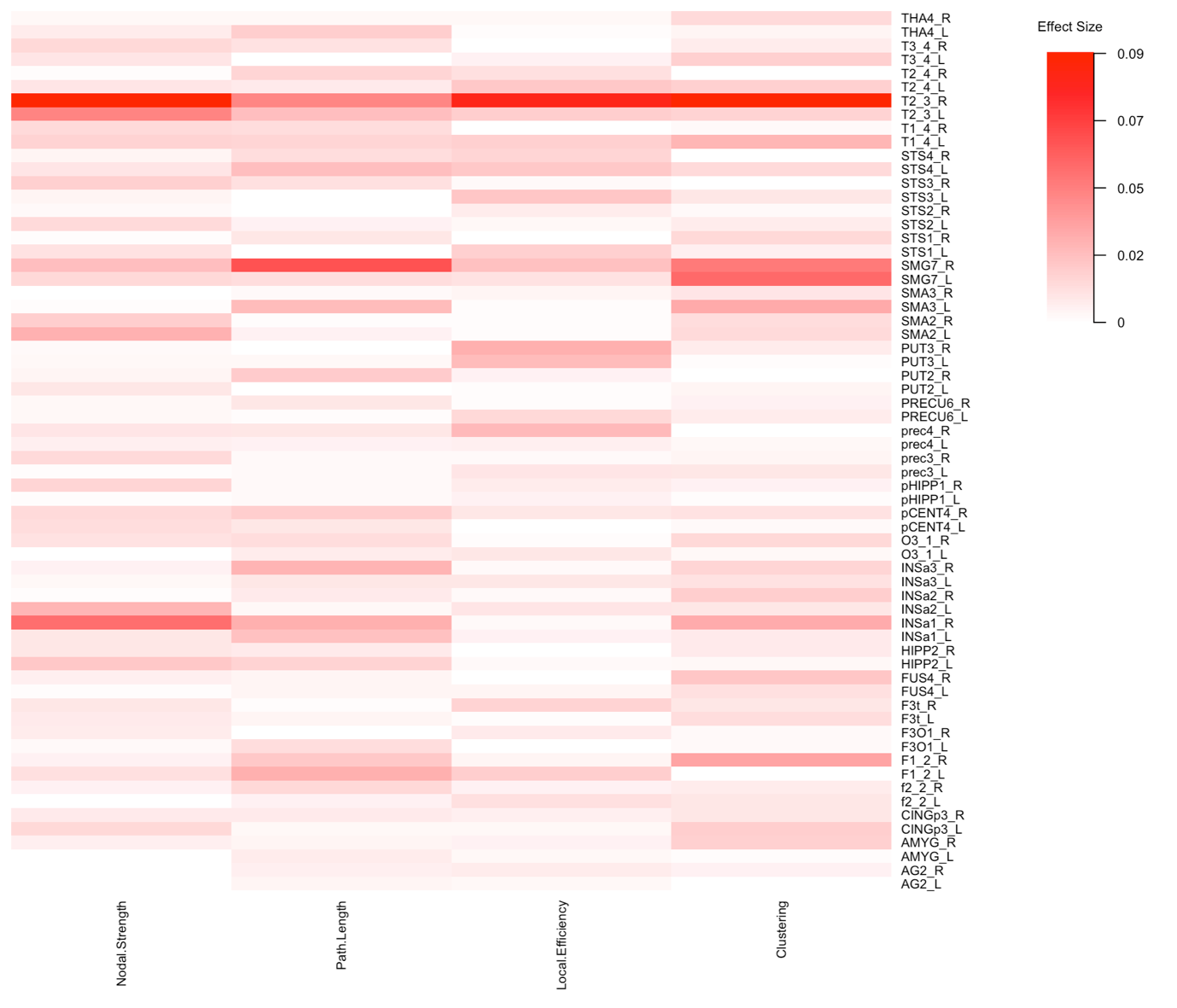
**Supplementary Figure 1. Heatmap depicting the effect sizes of metrics across nodes for different groups.** The effect sizes, representing the magnitude of differences between groups, are shown using a colour scale ranging from white (indicating no effect) to red (indicating small effects). No significant differences were observed between the groups, as indicated by the small effect sizes (maximum value of 0.09) across all metrics. The x-axis represents the metrics, and the y-axis lists the nodes.
